# Phytochrome A is required for light-inhibited germination of *Aethionema arabicum* seed

**DOI:** 10.1101/2025.02.26.640300

**Authors:** Zsuzsanna Mérai, Fei Xu, Anita Hajdu, László Kozma-Bognár, Liam Dolan

## Abstract

The germination of most seeds is influenced by the duration, intensity, and quality of light. The seeds of the model plant Arabidopsis are positive photoblastic and require light to germinate. The germination of negative photoblastic seeds is inhibited by white light. The molecular mechanisms that regulate negative photoblastic germination are unknown due to the lack of a suitable model plant.
We identified an accession with negative photoblastic germination in *Aethionema arabicum* that grows in semi-arid natural habitats. In a forward genetic screen, we identified a mutant – *koyash2* (*koy2*) – that is defective in negative photoblastic germination. There is a nonsense mutation in the gene encoding the phytochrome A photoreceptor in the *koy2* mutant.
Here we show that phytochrome A is required for negative photoblastic germination. The defective negative photoblastic phenotype of the *koy2* mutant is the result of defective inhibition of germination by the phytochrome A mediated high-irradiance response.
This is the first example of phytochrome A-mediated response controlling negative photoblastic seed germination in white, red, far-red, and blue light. We speculate that genetically encoded variation in phytochrome A-mediated germination responses is responsible for local adaptation of *Ae. arabicum* throughout the Irano-Turanian region.

**One sentence summary:** Identification and characterization of a phytochrome A null mutant demonstrates an active role of phytochrome A in light-inhibited seed germination in Aethionema arabicum, a negative photoblastic Mediterranean plant.

## Introduction

Natural selection acts on genetic variation that arises in wild populations. This genetic variation can control the development of diverse life history traits that may be adaptive. *Aethionema arabicum* (*Brassicaceae*) is an annual plant that grows in the Mediterranean ecosystems of Southeastern Europe and the Irano-Turanian region. There is considerable variation in life history traits among *Aethionema* species and different *Ae. arabicum* accessions isolated from different parts of its natural range. For example, life cycles may be annual or perennial (Mohammadin *et al*., 2017). Fruits may be dimorphic and the ratio between the two morphs varies between closely related *Aethionema* species and between different *Ae. arabicum* accessions (Lenser *et al*., 2016; Mohammadin *et al*., 2017; Arshad *et al*., 2019; Mérai *et al*., 2023). The dimorphic fruit morph ratio is modulated by environmental factors and impacts patterns of seed dispersal (Lenser *et al*., 2018). These variations in life history are genetically controlled and therefore likely adaptations to local environments.

Seed germination characteristics impact the life history of plants, and while they differ between species, there is also variation within species. White light modulates seed germination; positively photoblastic seeds require a light pulse for germination, while white light represses germination in negatively photoblastic seeds. Light-neutral seeds germinate in both darkness and light. Photoblastic responses are white light-specific and do not include wavelength-specific seed germination responses (Takaki, 2001). An *Ae. arabicum* accession from Turkey germinates in all light conditions (light-neutral) upon imbibition (Mérai *et al*., 2019). However, other accessions of *Ae. arabicum* are negatively photoblastic and darkness stimulates germination, while white light represses germination (Mérai *et al*., 2019). This variation may reflect adaptations to different environments. The negatively photoblastic trait is likely adaptive and could hypothetically promote germination in conditions where the seed is not directly exposed to light. Seedlings would be less susceptible to the desiccation caused by direct exposure to the sun during early stages of establishment when root growth occurs but before photosynthesis is necessary. The germination of negatively photoblastic seeds may be modulated by day length (Mérai *et al*., 2019, 2023, 2024). For example, the seeds of a negatively photoblastic accession from Cyprus (CYP) do not germinate in white light in long day conditions. However, seeds of the CYP accession germinate in white light in short day conditions. This promotes the germination and seedling establishment during the spring season when conditions are optimal for germination and seedling establishment, whereas it inhibits germination during the summer period when the conditions are deleterious for seedling establishment (Mérai *et al*., 2023, 2024). This indicates that some negatively photoblastic seeds can sense differences in day length to repress germination in one condition and promote germination in another.

The mechanism that promotes seed germination by light– positive photoblasty – is well characterized in a variety of species including *Arabidopsis thaliana* (*Brassicaceae*). In *A. thaliana*, light stimulates the production of gibberellic acid (GA) and reduces the level of the germination repressor, abscisic acid (ABA), in seed. Light is perceived in seed by the phytochrome photoreceptors and increases the expression of genes encoding enzymes that promote GA synthesis (*GA3OX*) and represses the expression of genes encoding enzymes that metabolize GA (*GA2OX*) (Seo *et al*., 2006). Light also reduces the expression of genes encoding proteins involved in ABA synthesis (*NCED5, NCED6*). Light signalling modulates the balance between the germination-promoting activity of GA and the germination-repressing activity of ABA (Seo *et al*., 2009).

Phytochromes are red and far-red absorbing photoreceptors in plants that exist in interconvertible forms; the red-light absorbing Pr (inactive) and the far-red light absorbing Pfr (active) form. Multiple genes encode phytochromes in most angiosperms. For example, there are five different phytochrome proteins – phyA, phyB, phyC, phyD, phyE – in *A. thaliana*. phyA and phyB are the dominant photoreceptors that control seed germination in *A. thaliana*. However, phyE is also involved and its function can be observed in *phy1 phyb* double mutants (Hennig *et al*., 2002).

While the absorption spectra of all *A. thaliana* phytochromes are identical, the specificity in light responses can be explained by the difference in the kinetics of the interconversion between the Pr and Pfr forms, cellular localization, and specific molecular interactions. The active Pfr form of phyA is light-labile and imported into the nucleus upon illumination and subsequently degraded (Somers et al., 1991). PhyB-E on the other hand are light-stable. The physiological responses are initiated by phyA, in low levels of Pfr that occur when plants are grown in high levels of far-red light or in low intensities of white light. Under these conditions, the active nuclear pool of phyA can be maintained by the constant nuclear import of a small Pfr amount that replaces the degraded fraction (Casal, Candia and Sellaro, 2014).

Phytochrome acts in discrete responses depending on the fluence level and wavelength of light (Casal, Sánchez and Botto, 1998). The very low fluence response (VLFR) regulates germination by the phy A photoreceptor (Shinomura *et al*., 1996). VLFR is active when the Pr:Pfr ratio is high and the relative levels of Pfr are very low. For example, VLFR-activated germination is induced by red light between 1-100 nmol m^-2^ total fluence or low intensity far-red light between 0.5 and 10 μmol m^-2^ total fluence in which as little as 0.01% of phytochrome is in the active Pfr form (Shinomura *et al*., 1996). The low fluence response (LFR) photoreversibly regulates germination at light intensities between 10-1000 μmol m^-2^ total fluence through the phyB photoreceptor; red light induces, and far-red light inhibits germination (Shinomura *et al*., 1996). The high intensity response (HIR) regulates hypocotyl and stem elongation and anthocyanin production but does not regulate seed germination of *A. thaliana* (Casal and Sánchez, 1998). The high irradiance response requires *circa* 60 μmol m^-2^ s^-1^ I and is mediated by phyA (Shinomura, Uchida and Furuya, 2000). Germination of *Ae. arabicum* seed is promoted by the VLFR and inhibited by the HIR but it is unknown if the low fluence response (LFR) also controls germination (Mérai *et al*., 2023). Identifying which phytochrome is involved in conferring germination and through which mechanisms it operates is an outstanding question in *Ae. arabicum* seed biology.

Phytochrome A and B promote germination in positively photoblastic *A. thaliana* seed (Shinomura *et al*., 1994; Lee *et al*., 2012). Less is known about the mechanism of light-repressed seed germination in negatively photoblastic plants such as the negatively photoblastic accessions of *Ae. arabicum*. Although it is known that phytochrome is required for negative photoblasty, the identity of the phytochrome(s) that is active during negative photoblasty in *Ae. Aethionema* is unknown (Mérai *et al*., 2023). To identify genes required for negatively photoblastic seed germination in *Ae. arabicum*, we carried out a genetic screen in the Cyprus (CYP) accession that is negatively photoblastic: it germinates only in the dark but does not germinate in the light. We screened for mutants that germinated in the light, reasoning that the genes required for light-repressed germination would be defective in these mutants. We identified *Koyash* (*koy2*) mutants that germinated in the light and carried a mutation in the phytochrome A gene. We show that phyA (KOY2) is required for light-induced repression of germination in this negatively photoblastic accession. However, phyA activity is required for two antagonistic responses in *Ae. arabicum* (CYP) seed. It promotes light-induced germination through VLFR while it represses germination in red and far-red HIR. These data indicate that the same photoreceptor is active in promoting or repressing germination in *Ae. arabicum* through the VLFR and HIR, respectively. These data also suggest that the divergence of negative photoblasty from a positive photoblastic state in *A. arabicum* involved regulatory changes downstream of the photoreceptor.

## Materials and Methods

### Plant and seed material

Experiments were conducted with *Aethionema arabicum* (L.) Andrz. ex DC. accessions CYP (obtained from Eric Schranz, Wageningen). Wild type and *koy2* plants were propagated for seed material under 16 h light/19°C and 8 h dark/16°C diurnal cycles, under ∼300 μmol m^-2^ s^-1^ light intensity. Wild type and *koy2* plants were randomly distributed on the shelves. Seeds were harvested upon full maturation and stored in darkness at 50% humidity and 24°C for at least three months, except for the experiments in Figure 4B and 4C where semi-dormant seed batches eight weeks after harvest were used.

### Plant chambers and light source

The mutant screen and white light germination assays were carried out in a Percival plant growth chamber equipped with fluorescent white light tubes (Philips). Red, far-red, and blue light applications were performed using customer-designed LED light sources, using LEDs from OptoSupply (www.optosupply.cn). The spectral properties and the LED types are as in (Mérai *et al*., 2023). Light spectra and intensity were measured by LED meter MK350S (UPRtek).

### *Aethionema arabicum* genome and annotations

*Aethionema arabicum* genome version 3.0, gene models, cDNA, and protein annotations (version 3.1) were obtained from *Ae. arabicum* database (https://plantcode.cup.uni-freiburg.de/aetar_db/) (Fernandez-Pozo *et al*., 2021). Gene accession numbers used in this study are listed in Supplementary Table S1.

### Forward genetic screening

For genetic screening, mutant seed bank and screening conditions were used as described in (Mérai *et al*., 2023). 20-30 seeds each from M2 or M3 seed batches were tested from approximately 1320 lines originating from independent mutagenized seeds. Seed germination assays were carried out in Petri dishes with wet filter paper at 14°C in a Percival growth chamber. Seeds were kept under continuous 160 μmol m^-2^ s^-1^ white light and scored for germination after 7 days. The germinating seeds were kept and propagated. One selected line with a stable mutant phenotype in two following generation was named as *koy2*. The *koy2* mutant line was backcrossed with the wild type and the segregation of the germination phenotype was determined in the progeny. As the original mutant line is hyposensitive and germinates with ∼80% at 160 μmol m^-2^ s^-1^, in contrast to 0% of the wild type, we expected the germination rate in the F2 backcross generation in case of a recessive mutation to be lower than 25%. The observed germination of F2 seed population under light was ∼17% and matched this expectation. Given the long hypocotyl phenotype in the *koy2* mutant under far-red light, the gene encoding PHYA (*Aa31LG1G5460*) was amplified with primers 5’ATGTCAGGAGCTAGGCCGAGTC and 5’TTTGTTTGCAGCTGCAAGTTCAG and sequenced throughout its entire full length. For genotyping the *koy2* mutant, the gene region was amplified using primers 5’TATAAGTTAGCTGCTAAAGCGA and 5’AGATGATCTTGGATGCATTCTTC, followed by Sanger sequencing of the amplicons.

### Germination assays

Germination assays were carried out in Petri dishes on wet filter paper using distilled water supplemented with 0.1 v/v % PPM™ (Plant Preservative Media, 5-Chloro-2-methyl-3(2H)-isothiazolone, Plant Cell Technology). 20-30 seeds were used for one replicate and three biological replicates were used from different seed batches. Germination in darkness was tested by transferring seeds onto wet plates in complete darkness without green safety light and subsequent wrapping the plates in two layers of aluminum foil and placing them in a dark box. All germination assays were done at 14°C and scored after 7 days. For light pulse experiments, 100 μmol m^-2^ s^-1^ red light and 10 μmol m^-2^ s^-1^ far-red light was used for 10 min (Figure 4B and C).

### Hypocotyl assay

Hypocotyl assays were carried out as described in (Mérai *et al*., 2023).

### RNA extraction and quantitative RT-PCR

To obtain RNA samples, wild type and *koy2* seeds were exposed to continuous light exposure for 24 h or kept in darkness. 80 μmol m^-2^ s^-1^ red light, 1 μmol m^-2^ s^-1^ far-red light, or 100 μmol m^-2^ s^-1^ blue light was used. Only seeds with intact seed coat were collected under safety green light. RNA extraction, cDNA synthesis, and RT-qPCR were performed as described (Mérai *et al*., 2019, 2023), using the primer pairs listed in Supplemental Table S1. The geometric mean of *Ae. arabicum* putative orthologues of *POLYUBIQUITIN10* (*AearUBQ10, Aa3LG9G835*) and *ANAPHASE-PROMOTING COMPLEX2* (*AearAPC2, Aa31LG10G13720*) was used for normalization. For each gene, the expression levels were presented as fold change relative to the level of the dark samples in wild type seeds, where the average expression was set to one. Statistical analysis was done using the SATQPCR tool (Rancurel *et al*., 2019). Error bars represent standard deviation of three biological replicates. Asterisks indicate significant differences from the wild type or *koy2* dark level with P-values as *P < 0.05, **P < 0.01, and ***P < 0.001 calculated with the Tukey test.

### Western blotting

To obtain protein samples, wild type and *koy*2 seeds were germinated in darkness at 14°C for 6 days. Seedlings were transferred under red (100 μmol m^-2^ s^-1^), far-red (1 μmol m^-2^ s^-1^), or blue (100 μmol m^-2^ s^-1^) illumination for 24 h or kept in darkness. For positive control samples, *A. thaliana* wild type Wassilewskija ecotype seeds were exposed to 4 h light to induce germination, followed by 6 days darkness at 22°C. Total protein extraction and Western blot analysis were done as described (Kevei *et al*., 2007). 50 μg protein per lane was analyzed. For the detection of phyA proteins, a primary polyclonal rabbit antibody (Wolf *et al*., 2011), 1:2400 dilution) and a horse radish peroxidase(HRP)-conjugated secondary anti-rabbit antibody (Dako, #: P039901-2, 1:3000 dilution) wwere used. Actin levels were determined using a monoclonal anti-actin primary antibody (Sigma, #: A0480, clone 10-B3, 1:10000 dilution) and an HRP-conjugated secondary anti-mouse antibody (Thermo Fisher Scientific, #: 31431, 1:10000 dilution). Chemiluminescent signals were detected and quantified as described (Hajdu *et al*., 2018).

## Results

### *koy2* mutant seeds are hyposensitive for light-inhibited germination

Seed of the Cyprus accession of *Aethionema arabicum* germinate in darkness, while increasing light intensity gradually leads to complete inhibition of germination (Mérai *et al*., 2019, 2023). We performed a forward genetic screen to identify mutants that germinate despite light exposure, conditions where wild type seeds are fully inhibited (Mérai *et al*., 2023). 20-30 seeds of each mutant line generated by γ-radiation were plated under 120-160 μmol m^-2^ s^-1^ continuous white light and germination was scored after 6 days. Mutated lines with germinating seeds were kept and propagated for two generations to validate the stable inheritance of the phenotype. 100% of the M3 seed generation of one line germinated at 96 μmol m^-2^ s^-1^ compared to 12.5% germination of the wild type (Figure 1A, B). Exposure to higher light intensities (166 and 213 μmol m^-2^ s^-1^) resulted in complete inhibition of germination in wild type. However, 71.6% and 8% of the mutant seed germinated in 166 and 213 μmol m^-2^ s^-1^ light, respectively (Figure 1A). We concluded that seeds of this line are hyposensitive for the inhibition of germination by light. The line was named as *koyash2* (*koy2*) after the god of sun in Turkic mythology.

**Figure 1.**
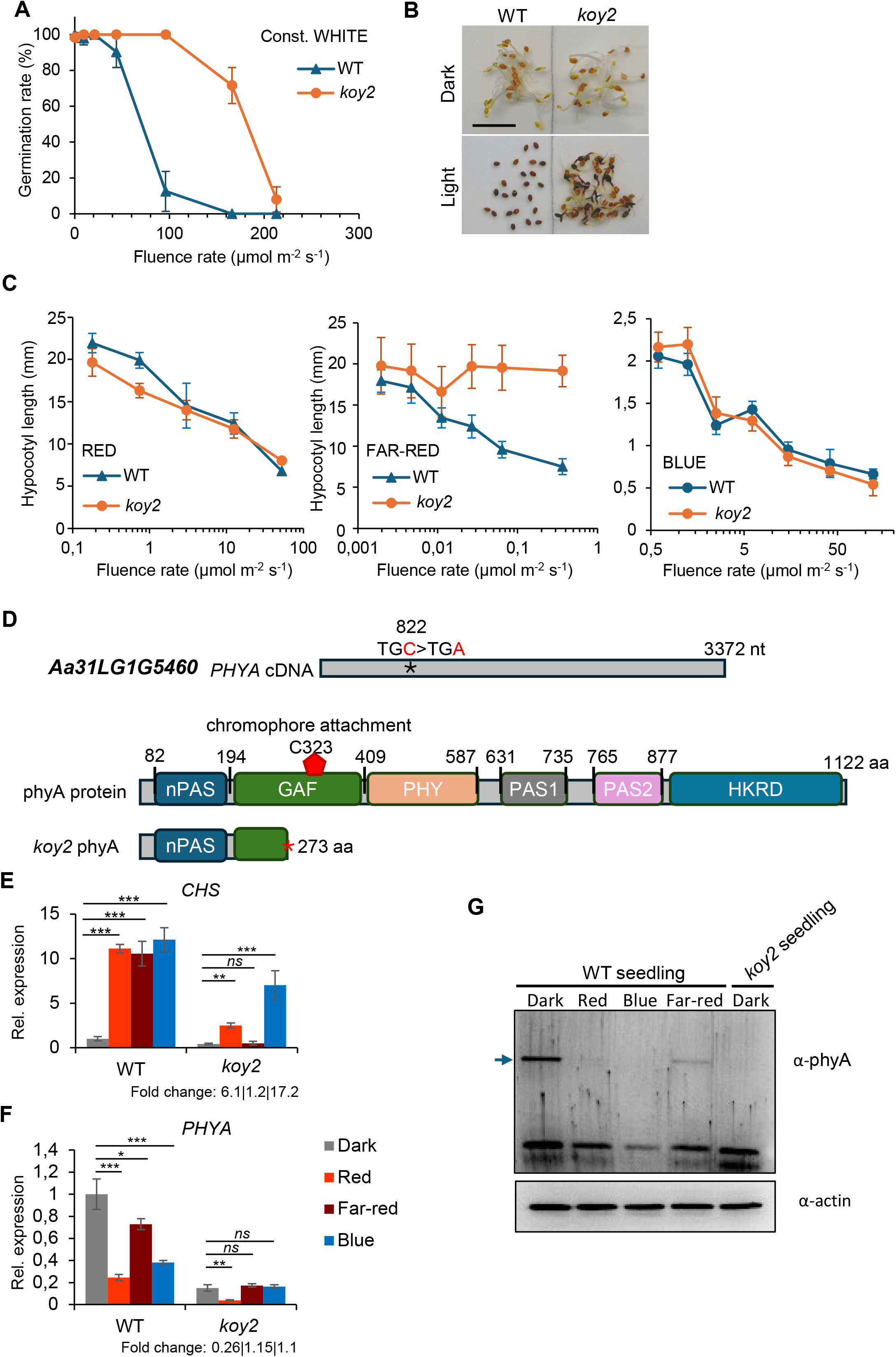
Identification of *koy2* as a *phya* null mutant. **(A)** Germination response of wild type (WT) and *koy2* mutant seeds under darkness (at light intensity 0) and continuous white light. Maximal germination was scored after 7 days. Error bars indicate the standard deviation of three biological replicates. **(B)** Images of wild type (WT) and *koy2* seeds after being kept in darkness or under 96 μmol m^-2^ s^-1^ white light for 7 days. **(C)** Hypocotyl elongation test of wild type (WT) and *koy2* seedlings. Seeds were kept in darkness for two days to induce germination, then illuminated with red, far-red, or blue light for 5 days during seedling growth. At least 12 hypocotyls were measured for each point. Error bars indicate the standard deviation. **(D)** Identified mutation in the coding sequence of the gene encoding phyA (*Aa31LG1G5460*). Protein domain structure of wild type phyA protein and the truncated protein in *koy2* lacking the chromophore binding site. **(E-F)** Relative expression of *AearCHS* (E) and *AearPHYA* (F) in wild type (WT) and *koy2* seeds, incubated in darkness, under red, far-red, or blue light for 24 h. Expression level is normalized to the level in wild type (WT) in darkness, which is set to 1. ‘Fold change’ numbers indicate the fold change of expression in red, far-red, or blue exposure of the *koy2* mutant relative to the level in *koy2* seeds incubated in darkness. Error bars indicate the standard deviation of three biological replicates. Asterisks indicate significant differences from the wild type or *koy2* dark level with P-values as *P < 0.05, **P < 0.01, and ***P < 0.001 calculated with the Tukey test. **(G)** Accumulation of phyA protein (indicated with arrow) in wild type (WT) and *koy2* seedlings.

The hyposensitive phenotype of *koy2* seeds suggested that the mutation affects either the light perception or the phytohormone metabolism. To discriminate between these alternatives, we compared hypocotyl elongation under monochromatic red, far-red, and blue light in wild type and *koy2* mutants. We hypothesized that if the mutation affects light perception, the hypocotyl shortening response of the mutant seedlings will significantly differ from the wild type shortening response. Seeds were plated and kept in darkness for two days to induce germination. Germinating seeds were then illuminated with increasing light intensities of red, far-red, and blue light, for five days. The hypocotyl shortening response was similar in the *koy2* mutant and the wild type under red and blue light; both *koy2* and wild type elongated less in increasing light (Figure 1C). This indicates that hypocotyl elongation in red and blue light is not defective in the *koy2* mutant. However, the length of *koy2* seedlings in far-red light was the same as in darkness, indicating that the *koy2* mutant is non-responsive to far-red light (Figure 1C).

In angiosperms, phytochrome A is the far-red photoreceptor; complete loss-of-function mutants of Arabidopsis *PHYA* develops long hypocotyls under far-red light. Therefore, we hypothesized that *koy2* is a *phya* mutant in Aethionema. The sequencing of *PHYA* cDNA in *koy2* revealed a single C to A nucleotide change at the position of 822 nucleotides, resulting in a predicted premature termination codon after amino acid 273 (Figure 1D). Seedlings with long hypocotyls grown under far-red light were homozygous for the *koy2* mutant allele while seedlings with short hypocotyls were heterozygotes or carried the wild-type *PHYA* allele (Supplementary Table S2). The predicted truncated phyA protein lacks the chromophore attachment site that is essential for light perception. Therefore, we conclude that *koy2* is a complete loss of function *phya* mutant allele (Figure 1D). To test this hypothesis, we measured the steady state levels of *CHALCONE SYNTHASE (CHS)* mRNA that accumulates at higher steady state levels in seeds and seedlings upon red, blue and far-red light, than in the dark (Mérai *et al*., 2023). In wild-type seeds, CHS mRNA levels were higher in red, far-red, and blue light than in dark-grown controls. However, steady state CHS mRNA levels in *koy2* seeds were identical in far-red light compared to those kept in the dark. Furthermore, the steady state CHS mRNA levels in *koy2* seeds were significantly higher in red and blue light than in darkness, as in wild type. These data indicate that the induction of *CHS* expression by far-red light is defective in *koy2* while *CHS* induction in red and blue light is independent of *KOY2* function. These data are consistent with the hypothesis that KOY2/phyA are required for the far-red responses in *Ae. arabicum*.

Light reduces steady state levels of *PHYA* mRNA and promotes phyA protein degradation by the 26S proteosome (Somers *et al*., 1991; Cantón and Quail, 1999). To test if the transcriptional and post-translational repression of PhyA also occurs in the wild type and the *koy2* mutant of *Ae. arabicum*, we measured levels of PhyA mRNA and protein. Steady state levels of *PHYA* mRNA level were highest in seeds incubated in darkness and significantly lower in red, blue, or far-red light-illuminated wild-type seeds (Figure 1F). The mRNA level of *koy2* mutant *PHYA* in the darkness was 15% the levels of wild type (Figure 1F).

*PHYA* mRNA levels were lower in red light in *koy2* than in darkness, indicating that the phyB-mediated repression of *PHYA* gene expression is functional in the *koy2* seed (Figure 1F).

The *PHYA* mRNA level did not change significantly in far-red light in *koy2* seed, further confirming that *koy2* is non-responsive to far-red light. The *PHYA* mRNA level was unchanged in *koy2* in blue light, suggesting the autorepression of *PHYA* expression under blue light (Figure 1F).

phyA protein is a light-labile photoreceptor quickly degraded in all light conditions in *A. thaliana*. We set out to test if phyA is light-labile in *Ae. arabicum*. phyA protein was detectable in dark-grown wild type *Ae. arabicum* seedlings. However, the protein was undetectable after 4 days of constant red or blue illumination and less abundant under far-red light than in darkness (Figure 1G). These data demonstrate that phyA protein is degraded upon exposure to light. Furthermore, phyA protein was not detected in the *koy2* mutant seedlings in darkness, indicating that phytochrome protein does not accumulate in the *koy2* mutant (Figure 1G).

Taken together, these data indicate that the *koy2* mutant is a *phya* complete loss of function mutant that is hyposensitive to the light-inhibited germination of *Ae. arabicum* seed.

### Loss-of-function *phyA (koy2)* mutant seeds germinate in red, far-red, and blue light

Red, blue, and far-red light inhibit germination of wild type *Ae. arabicum* in an intensity-dependent manner; higher light intensities lead to progressive reduction in germination (Mérai *et al*., 2019, 2023). To determine the wavelength-specific action of phyA in germination inhibition, the germination of wild type and *koy2* seed germination was measured under increasing intensities of red, far-red, and blue light illumination. 100% of *koy2* seeds germinated under 300 μmol m^2^ s^1^ red light, while no wild type seed germinated at 162 μmol m^2^ s^1^ and above (Figure 2B). These data are consistent with the hypothesis that red light repression of germination requires phyA (KOY2) activity. Over 95% of *koy2* seeds germinated in far-red light, while wild type did not germinate, even at very low light intensities (0.03 μmol m^2^ s^1^) (Figure 2A), consistent with the hypothesis that far-red light repression of germination requires phyA (KOY2) activity. *koy2* seed germination was hyposensitive for blue light; 72% of *koy2* mutant seed germinated in blue light while 0% of wild type seed germinated at 122 μmol m^2^ s^1^ (Figure 2C), consistent with the hypothesis that blue light repression of germination requires phyA (KOY2) activity. These results indicate that phyA mediates the inhibition of seed germination by red, far-red, and blue light in *Ae. arabicum*.

**Figure 2.**
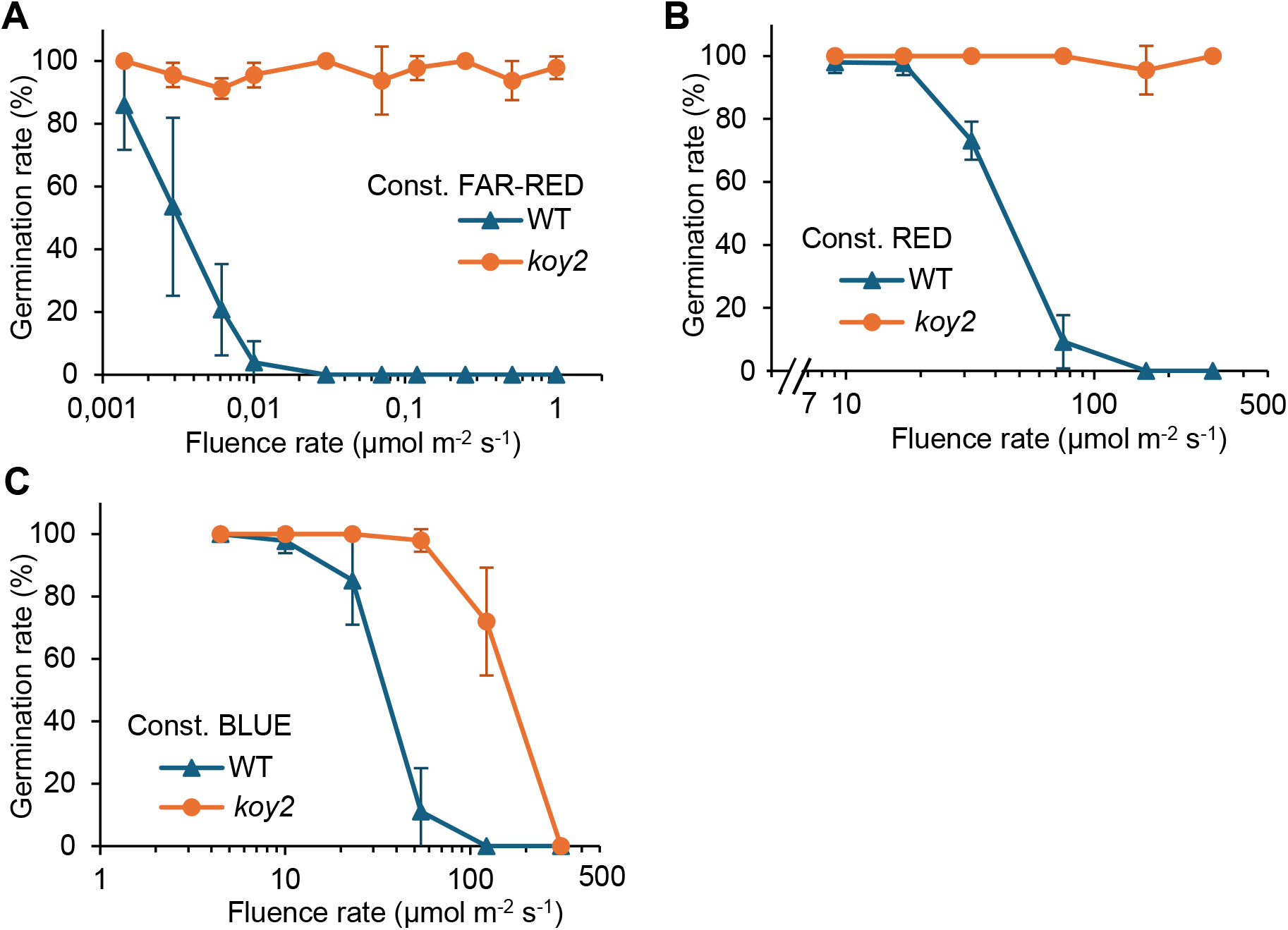
Germination response of wild-type (WT) and *koy2* mutant seeds under far-red (A), red (B), and blue (C) light. Maximal germination was scored after 7 days. Error bars indicate the standard deviation of three biological replicates.

### phyA (KOY2) promotes the expression of genes encoding proteins that reduce GA levels and increase ABA levels in seeds in red, far-red, and blue light

#### Red light

Red light represses *Ae. arabicum* seed germination (Mérai *et al*., 2019, 2023). Phytochromes regulate germination through a signalling cascade that leads to the transcriptional regulation of the genes encoding enzymes of GA and ABA metabolism (Seo *et al*., 2006). To test the role of phyA (KOY2) in GA metabolism during the repression of germination by red light, we compared the steady state level of mRNAs for key enzymes for GA synthesis (*AearGA3ox1*) and GA degradation (*AearGA2ox3*). Red light-exposure of wild-type plants increased the steady state levels of *AearGA2ox3* 50-fold compared to dark-treated wild type. Since AearGA2OX3 catabolizes GA, the higher levels of *AearGA2ox3* mRNA levels likely reduce GA levels and contribute to the repression of germination by light (Figure 3). The light-induced increase in steady state levels of *AearGA2ox3* mRNA was much lower in *koy2* mutants than in wild type seed; red light increased steady state levels 4.8 times in *koy2* compared to 50 times in the wild type. These data demonstrate that phyA (KOY2) is required for the increase in expression of *AearGA2ox3* during red light-mediated repression of germination. To test if phyA (KOY2)-mediated red light signalling represses *AearGA3ox1* expression, the steady state levels of *AearGA3ox1*mRNAs were compared in wild type and *koy2* mutants. In wild type, steady state levels of *AearGA3ox1* mRNA were lower in red light-treated plants than in darkness, consistent with the hypothesis that red light represses germination through repression of GA synthesis. Furthermore, steady state levels of *AearGA3ox1* mRNA were higher in *koy2* mutants than in wild type. These data suggest that red light repression of *AearGA3ox1* is phytochrome dependent. Taken together, these data suggest that phyA (KOY2)-mediated red light signalling promotes the expression of AearGA2OX3, which catabolises GA and represses the expression of AearGA3OX1, required for GA synthesis, during red light-induced repression of germination in *A. arabicum* seed.

**Figure 3.**
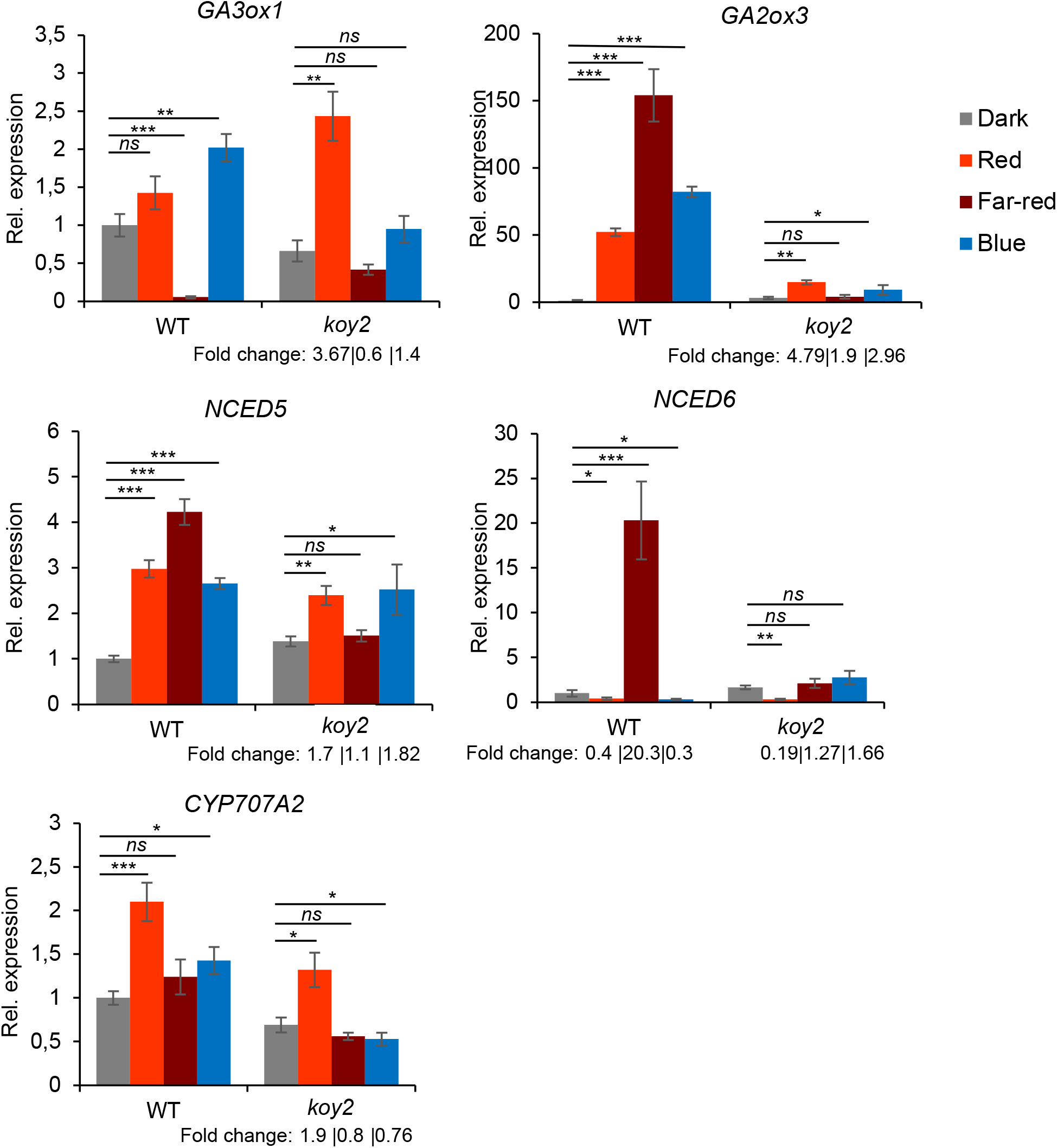
Light-induced gene expression of genes encoding GA and ABA synthesis and catabolic enzymes. Relative expression of selected genes in wild type (WT) and *koy2* seeds incubated in darkness, or seeds incubated under 80 μmol m^-2^ s^-1^ red light, 1 μmol m^-2^ s^-1^ far-red light, or 100 μmol m^-2^ s^-1^ blue light for 24 h. Expression level is normalized to the level in wild type (WT) in darkness, which is set to 1. ‘Fold change’ numbers indicate the fold change of expression in red, far-red, or blue exposure of the *koy2* mutant relative to the level in *koy2* seeds incubated in darkness. Error bars indicate the standard deviation of three biological replicates. Asterisks indicate significant differences from the wild type or *koy2* dark level with P-values as *P < 0.05, **P < 0.01, and ***P < 0.001 calculated with the Tukey test.

**Figure 4.**
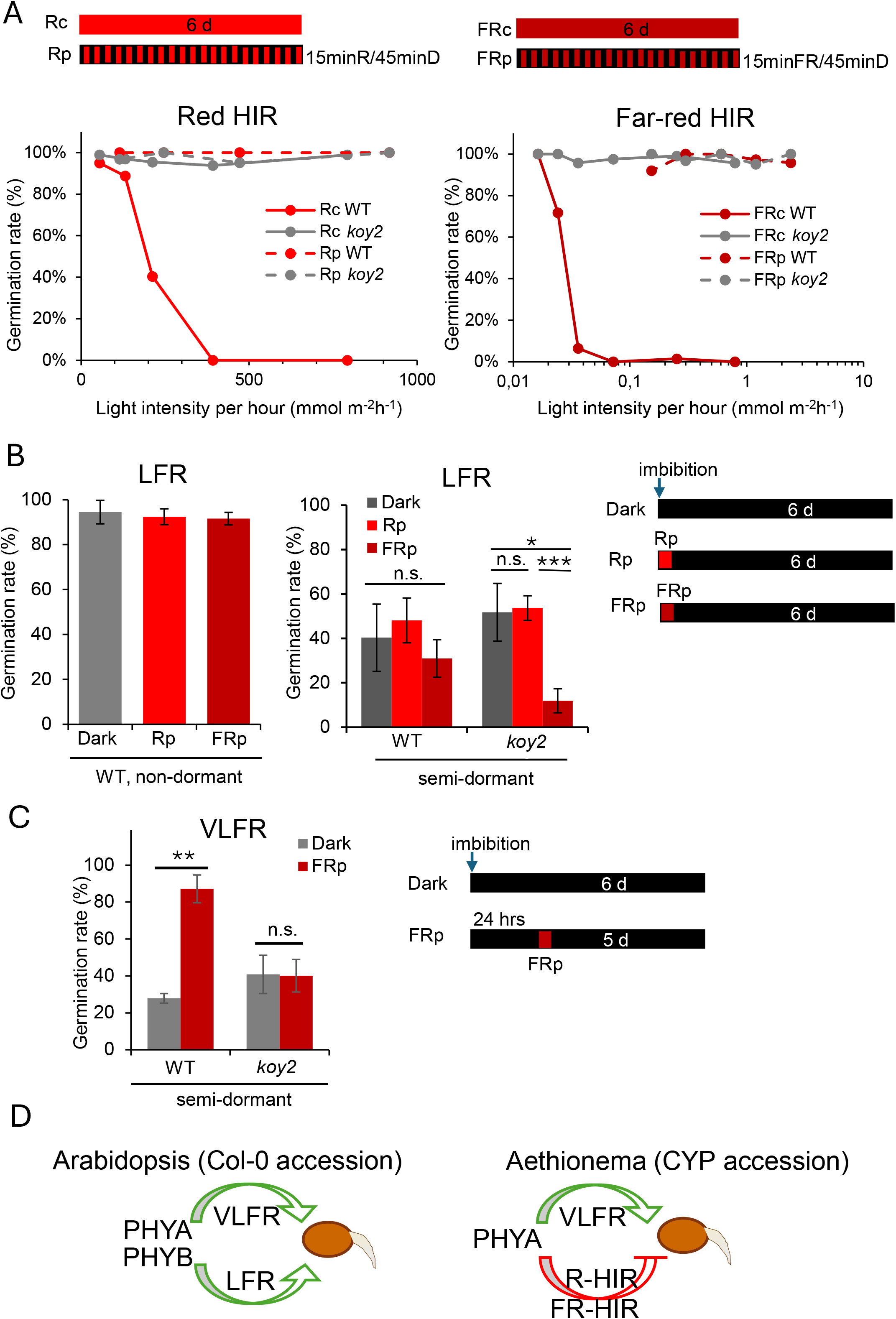
Phytochrome action modes in *Ae. arabicum* seeds. **(A)** Total germination of wild type and *koy2* seeds under constant red (Rc, left panel) or far-red light (FRc, right panel) or 15 min light pulses (Rp, dashed lines, left panel, or FRp, dashed lines, right panel) intermitted with 45 min dark periods. Dots indicate the average of three biological replicates. Maximal germination was scored after 6 days. **(B)** Total germination of wild type non-dormant (left panel) or semi-dormant (right panel) and *koy2* semi-dormant seeds. Seeds were exposed to 10 min of 100 μmol m^-2^ s^-1^ red light pulse (Rp) or 10 μmol m^-2^ s^-1^ far-red pulse (FRp) followed by 6 days incubation in darkness. ‘Dark’ indicates that seeds were kept in darkness without any light pulses. Error bars indicate the standard deviation of three biological replicates. **(C)** Total germination of semi-dormant wild type and *koy2* seeds. Seeds were imbibed in complete darkness. 24 h after imbibition, seeds were illuminated for 10 min with 10 μmol m^-2^ s^-1^ far-red (FRp) or kept in darkness. Germination was scored 6 days after the imbibition. (B-C) Asterisks indicate significant differences in pairwise comparisons within one genotype with P-values as *P < 0.05, **P < 0.01, and ***P < 0.001 calculated with the Students *t*-test. **(D)** schematic representation of the phytochrome action modes in the positive photoblastic Arabidopsis seeds and the negative photoblastic seeds in *Ae. arabicum* (CYP accession). Green arrows indicate germination induction and red arrows indicate germination inhibition.

ABA represses seed germination (Ali *et al*., 2022). To test the role of phyA (KOY2) in ABA-mediated repression of germination, we measured the steady state levels of mRNAs for proteins involved in ABA synthesis, *AearNCED5, AearNCED6*, and in ABA degradation (*AearCYP707A2*) in red light. *AearNCED5* and *AearCYP707A2 mRNA* levels were higher in red light than in darkness, while *AearNCED6* level was lower in red light compared to darkness. This higher level of *AearNCED5* in red light-treated seeds is consistent with the hypothesis that red light stimulates the production of ABA. Furthermore, *AearNCED5, AearNCED6*, and *AearCYP707A2* mRNA levels were almost identical in wild type and *koy2* under red light. These data indicate that the levels of genes encoding enzymes for ABA metabolism do not require phyA (KOY2) activity.

The data suggest that phyA (KOY2) represses GA synthesis and induces GA degradation during red light-mediated repression of germination.

#### Far-red light

Far-red light represses *Ae. arabicum* germination (Mérai *et al*., 2019, 2023). To determine the effect of far-red light on the expression of genes that promote germination (*AearGA3ox1*) and repress germination (*AearGA2ox3, AearNCED5*, and *AearNCED6*), we compared the corresponding steady state mRNA levels in seed grown in far-red light and darkness. In wild type, steady state levels of *AearGA3ox1* were lower in far-red light than in darkness while steady state levels of *AearGA2ox3, AearNCED5*, and *AearNCED6* were higher in far-red light than in darkness (Figure 3). These data suggest that far-red light represses GA synthesis and promotes GA degradation and ABA synthesis (Merai et al., 2023). However, the steady state levels of *AearGA3ox1, AearGA2ox3, AearNCED5*, and *AearNCED6* mRNAs in far-red light and darkness in the *koy2* mutant were indistinguishable, indicating that phyA (KOY2) is required for their expressional changes in far-red light. These data indicate that phyA is required for the repression of *AearGA3ox1* and induction of *AearGA2ox3, AearNCED5*, and *AearNCED6* in far-red light. Together this would reduce GA levels and increase ABA levels in far-red light compared to controls in the wild-type seeds.

#### Blue light

Blue light represses *Ae. arabicum* germination (Mérai *et al*., 2019, 2023). To test the function of phyA in blue light repression of germination, steady state mRNA levels of *AearGA3ox1, AearGA2ox3, AearNCED5, AearNCED6*, and *AearCYP707A2* were measured in wild type and *koy2* seed grown in 100 μmol m^2^ s^1^ blue illumination in which no wild type seed germinated and approximately 80% of *koy2* seeds germinated. Steady state levels of *AearGA3ox1, AearGA2ox3, AearNCED5*, and *AearCYP707A2* were higher in wild-type plants grown in blue light than in the dark. In *koy2* seed, the difference in mRNA levels of *AearGA2ox3* and *AearNCED*5 mRNAs between blue light-treated and dark-grown plants was much less than in the wild type. For example, in wild type, *AearGA2ox3* mRNA levels were 82-fold higher in blue light than in darkness but only 2.96-fold higher in blue light than in darkness in *koy2* mutant seed (Figure 3). These data indicate that phyA is required for the induction of the genes that repress germination – *AearGA2ox3* and *AearNCED5* – in blue light-treated *Ae. arabicum* seeds. The expression of *AearGA3ox1* and *AearCYP707A2* was identical between blue light-treated wild type and *koy2* mutants, indicating that their expression is not modulated by phyA. Taken together, the expression of negative regulators of seed germination – *AearGA2ox3* and *AearNCED5* – is induced in wild type in red, far-red, and blue light conditions. Their expression is lower in *koy2* mutant than in wild type in each of the light conditions. This indicates that phyA (KOY2) activity promotes the expression of the germination repressors AearGA2OX3 and AearNCED5 in red, far-red, and blue light in *Ae. arabicum*.

### phyA (KOY2) is required for the high irradiance response-repression of germination in non-dormant seed

*Ae. arabicum* seed germination does not require light, because 100% of ripened non-dormant seeds germinate in darkness. While the inhibitory effects of light on germination can be tested on after-ripened, non-dormant seed, the promotion of germination by light can be tested by evaluating the effects of light exposure on semi-dormant seed, a state that is reached if seeds are stored for 8 weeks after harvest.

*Ae. arabicum* seed germination is repressed by the high irradiance response (Mérai *et al*., 2023). To test if germination inhibition by high irradiance response is mediated by phyA, we compared the germination of wild type and *koy2* seeds under constant light where the high irradiance response is active and represses germination. Illumination with continuous red or far-red light inhibited germination of wild type seed. Zero % of wild type germinated when imbibed seed was illuminated in 392 mmol m^-2^ h^-1^ continuous red or 0.072 mmol m^-2^ h^-1^ continuous far-red light, confirming that the high irradiance response repressed germination (Figure 4A). By contrast, 95.7% of *koy2* seeds incubated in 917 mmol m^-2^ h^-1^ continuous red, and 100% *koy2* seed incubated in 0.6 mmol m^-2^ h^-1^ far-red light germinated (Figure 4A).

These data are consistent with the hypothesis that phyA (KOY2) is required for the high irradiance repression of germination.

A defining characteristic of the high irradiance response is the requirement for continuous illumination to trigger a physiological response; the high irradiance response is not induced if light application is interrupted by intermittent dark pulses (intermittent light). If the high irradiance response represses germination, we hypothesized that (i) the germination percentage of wild type would be higher in intermittent light than under continuous light and (ii) *koy2* mutants would be similar in intermittent light, because HIR would be inactive in both genotypes. With intermittent red (red light with intermittent darkness) or far-red (far-red with intermittent darkness) exposure, wild type seeds germinated (100% in red and 95% in far-red) while 0% of wild type seed grown in continuous red or far-red light germinated. Second, similar germination rates were found in wild type (100% in red and 95% in far-red) and *koy2* mutants (100% in red and far-red). These data are consistent with the hypothesis that the high irradiance response is defective in *koy2* mutants.

The defective germination of *koy2* mutants indicates that phyA (KOY2) activity is required for the repression of seed germination by the red and far-red high irradiance response in *Ae. arabicum*.

### PhyA is required for *VLFR-*promoted germination of semi-dormant seed

The very low fluence response promotes germination in *Ae. arabicum* (Mérai *et al*., 2023). Since VLFR promotes germination, it can be evaluated in semi-dormant seed 8 weeks after harvest. To test if phyA is required for the very low fluence response control of seed germination, we determined the phenotype of *koy2* mutants in conditions where a light pulse 24 h after imbibition activates this response. 28% of semi-dormant wild type seed germinated in the dark (optimal germination conditions) (Figure 4C). A far-red light pulse 24 h after imbibition increased the germination to 87%. This confirms that the very low fluence response promotes seed germination in wild type *Ae. arabicum* seed (Figure 4C) (Mérai *et al*., 2023). By contrast, *koy2* germination rate was the same in seed incubated in darkness (40%) and in darkness supplemented with a red light pulse after 24 h (40%) (Figure 4C). The lack of germination induction after a red light pulse at 24 h in *koy2* mutant seed indicates that phyA is required for the very low fluence response that promotes germination in semi dormant *Ae. arabicum* seed.

### The low fluence response does not control germination of semi dormant seed but phyA (KOY2) promotes germination after a pulse of far-red light

The role of the phytochrome-mediated low fluence response in germination in *Ae. arabicum* is unknown. We set out to test if the low fluence response controls germination. A defining characteristic of the low fluence response is reversibility; red light pulses induce the low fluence response while far-red pulse inactivates the response. We hypothesized that if the low fluence response controls germination, the germination rate of semi-dormant seed would be higher after a red pulse and lower after far-red pulse than in darkness.

First, we tested the effect of red and far-red pulses on non-dormant seed. Over 95% of after-ripened, non-dormant *Ae. arabicum* seeds germinated in darkness, indicating that light is not required for germination of seeds at this stage of maturity. Similar levels of germination were observed when a red pulse or far-red pulse was followed by darkness (Figure 4B). This indicates that a short light pulse does not impact germination in non-dormant seed (Figure 4B). These data suggest that the low fluence response does not operate in non-dormant seed to promote germination.

To determine if the low fluence response controls germination of semi-dormant *Ae. arabicum*, seed, we measured germination of seed batches approximately 8 weeks after harvest. The maximum seed germination ranged from 20 to 50% when wild-type seeds were plated in the dark (optimal) conditions. To test if pulses of red or far-red light impacted germination, semi-dormant seed were exposed to light pulses upon imbibition followed by dark incubation. The germination of wild type semi-dormant seeds was not significantly different in darkness and in applications where a red pulse was followed by darkness (Figure 4B). This indicates that the low fluence response does not control the germination of semi-dormant wild-type seed.

If the low fluence response does not control germination in *Ae. arabicum*, we hypothesized that the germination of semi-dormant seed would be identical in wild type and *koy2* (*phya)* mutants. To test this hypothesis, we compared the germination of wild type and *koy2* semi-dormant seed in conditions where the low fluence response would be active if present. The germination rates of wild type and *koy2* mutant seed was similar when incubated in darkness or after receiving a red pulse. This indicates that the low fluence response is not activated by red light in either wild type or the *koy2* mutant. These data further support the hypothesis that the low fluence response does not control seed germination in *Ae. arabicum*.

Next, we compared wild type semi-dormant seed germination in darkness with germination in darkness after a pulse of far-red light. 40% of wild type germinated in darkness while 31% germinated after a far-red light pulse. This indicates that far-red light at imbibition does not significantly affect germination in wild type semi-dormant seed. However, germination was significantly lower after a far-red pulse in the *koy2* mutant (11.9%) than in darkness (51.8%) (Figure 4B). These data indicate that phyA promotes germination induced by a pulse of far-red light at imbibition in semi-dormant seeds of *Ae. arabicum*.

Taken together, these data provide no evidence for the low fluence response in wild type seeds. However, phyA (KOY2) is required for the stimulation of germination by far-red light.

## Discussion

We report the discovery that phyA is required for negative photoblastic germination in *Ae. arabicum* (CYP accession) seeds. We identified a mutant *(koy2*) in *Ae. arabicum* (CYP accession) in a forward genetic screen that is defective in light-inhibited germination. *koy2* mutant seeds are less responsive to white and blue light than wild type, and unlike wild type do not respond to red and far-red light. The mutation in the *koy2* mutant encodes a premature stop codon in the gene encoding the phyA, the far-red photoreceptor in *Ae. arabicum*. The germination phenotype of the *koy2* mutant demonstrates that phyA is required for the repression of germination by white, red, far-red, and blue light in *Ae. arabicum*. This is the first detailed description of a role for phyA in the control of seed germination in a negatively photoblastic species, where white light strongly represses germination.

Phytochromes act through three distinct action modes, HIR, VLFR, and LFR. We demonstrate that both the HIR and VLFR operate during phyA-modulated germination in *Ae arabicum*. We show that phyA-mediated HIR promotes negative photoblasty in *Ae. arabicum*. Both red and far-red light-mediated HIR repress germination. While there are numerous examples of far-red light-mediated HIR in plants, this is only the second example of red light-induced HIR controlling germination. Previously, Appenroth (2006) demonstrated that red-light-induced HIR-repressed germination by ∼50% (Appenroth *et al*., 2006). In *Ae. arabicum*, the HIR effect is much stronger; 100% of seeds do not germinate when imbibed in red light HIR conditions.

HIR and VLFR function antagonistically in photoblastic seed germination in *Ae. arabicum*. Red and far-red HIR represses germination while far-red VLFR promotes germination. This antagonism between HIR and VLFR has also been demonstrated in the weed *Datura ferox* where far-red partially represses germination through HIR and induced germination through VLFR (Arana *et al*., 2007).

We demonstrate that phytochrome-mediated LFR does not act in *Ae. arabicum* to control seed germination. Phytochrome-mediated LFR controls germination in many species (Borthwick *et al*., 1952; Shropshire *et al*., 1961; Negbi and Koller, 1964). This action mode has been characterized in detail in *A. thaliana* and *Lactuca sativa* (lettuce) (Hartmann, 1966; Shinomura *et al*., 1996). In these species, red light promotes germination, and far-red light represses germination. This red light promotion and far-red light repression of germination is a characteristic of the LFR action mode and is mediated by phytochrome B, which is present in mature seed. Exposure to red light increases the Pfr:Pr ratio and activates the germination program through a PIF-dependent mechanism that increases GA levels and decreases ABA. Exposure to far-red light on the other hand, lowers the Pfr:Pr ratio and results in a decreased GA:ABA ratio. We demonstrated that the steady state mRNA level of the genes encoding the enzyme involved in GA degradation (*AearGA2ox3*) and ABA biosynthesis (*AearNCED5*) are much higher in both red and far-red light than in darkness. This would be expected to result in a decreased GA:ABA ratio, which would inhibit germination, in both red and far-red light. This is consistent with the hypothesis that red light does not promote germination through LFR in *Ae. arabicum*.

We demonstrated that phytochrome-mediated HIR repress germination and phytochrome-mediated LFR has no function in germination control in the CYP accession of *Ae. arabicum*. It has been proposed that the photoblastic mode – negative or positive – of seeds is determined by the HIR in negative photoblastic seed and LFR in positive photoblastic seed. There are many examples where LFR is involved in positive photoblastic seed germination (Shinomura *et al*., 1996; Appenroth *et al*., 2006). However, there are no examples where the presence of HIR and lack of LFR has been shown to be involved in negative photoblastic seed. Our demonstration that red and far-red light HIR are required for negative photoblastic germination supports this hypothesis. We provide the first complete set of evidence that supports the assertion that the distinct phytochrome action modes define seed germination behavior.

The photoblastic phenotype is an adaptive seed trait that has likely evolved as an adaptation for seedling establishment in the natural habitat. The Cyprus accession of *Ae. arabicum* grows in an exposed habitat with high illumination. Seedling establishment is poor in summer, when light levels and temperature are high and little water is available. Therefore, avoiding germination during the long days of summer may be adaptive. We speculate that phyA-mediated HIR represses germination during the summer in this accession of *Ae. arabicum*. Other accessions of *Ae. arabicum* are light neutral – seeds germinate in both darkness and light. We speculate that genetically encoded variations in phytochrome A-mediated light responses are responsible for local adaptation of *Ae. arabicum* throughout the Irano-Turanian region.

## Supporting information

Suppelemntal Table S2

## Acknowledgement

We thank Eric M. Schranz for providing Aethionema seed stocks. We also thank the staff of the Vienna BioCenter Core Facilities GmbH (VBCF), a member of Vienna BioCenter (VBC), Austria, especially the Plant Sciences Facility for the growth of the plants, the Molecular Biology Unit for providing multiple reagents, the Vienna Covid-19 Detection Initiative (VCDI) for generating a safe work environment during the pandemic and the VBC Child Care Center. We thank Nicole Lettner for the technical support. We thank Ortrun Mittelsten Scheid for the fruitful discussions and advices throughout the project.

## Author contributions

ZM planned and designed the research; ZM, FX, and AH performed the experiments; ZM, FX, LKB, and LD analyzed and interpreted the data; ZM and LD wrote the paper. All authors approved the submitted version.

## Supporting Information

**Supporting Table S1 List of primers used for quantitative RT-PCR analysis**.

**Supporting Table S2. Cosegregation of long hypocotyl phenotype under far-red with the mutation identified in *koy2***.

