## Supplemental Table S1 for "Phytochrome A is required for light-inhibited germination of *Aethionema arabicum* seed"

**Supporting Table S1 List of primers used for quantitative RT-PCR analysis.**

| **Name** | **Nucleotide sequence** | **Accession number v3.1** |
| --- | --- | --- |
| AearUBQ10_for | GAGGATGGCCGAACATTG | *Aa3LG9G835 (v3.0)* |
| AearUBQ10_rev | TGCCCGTTAGGGTTTTGA |  |
| AearAPC2_for | TCTCCTGCAATCGAGGACTT | *Aa31LG10G13720* |
| AearAPC2_rev | GCAGTGAGCAACCGGTATTT |  |
| AearCHS_for | GGCTCAAAGAGCTGATGGTC | *Aa31LG5G11220* |
| AearCHS_rev | TCGGTCATGTGGTCACTGTT |  |
| AearPHYA_for | GGAGAAGTCTTCGGGACACA | *Aa31LG1G5460* |
| AearPHYA_rev | TTTCTCCGGTTCTTGACTGG |  |
| AearGA3ox1_for | TCTTCGTCACCTCCCTGACT | *Aa31LG7G270* |
| AearGA3ox1_rev | GATGAGCGGGAGAGTTGTGT |  |
| AearGA2ox3_for | CGCGTCTCTCTTAACCCAAC | *Aa31LG10G6850* |
| AearGA2ox3_rev | TCACATGCCTTGACCATTTG |  |
| AearNCED5_for | GCCGTTTGATCTTGACGCTC | *Aa31LG6G5570* |
| AearNCED5_rev | ACGGAGTTTAGTTTACGGCGT |  |
| AearNCED6_for | GCTTCTTCAGCTCTCGACAA | *Aa31LG8G10550* |
| AearNCED6_rev | GAACCGTTGGATCAGTCGGT |  |
| AearCYP707A2_for | GCGGTTCCAACAAAGAAAAC | *Aa31LG5G7160* |
| AearCYP707A2_rev | GAGTGGCGAAGAAGGAATTG |  |
