## Supplementary material for "Phytochrome A is required for light-inhibited germination of *Aethionema arabicum* seed": Suppelemntal Table S2

**Supporting Table S2.** **Cosegregation of long hypocotyl phenotype under far-red with the mutation identified in *koy2*.**

| ♀ *koy-2* x ♂ WT | |  |  |
| --- | --- | --- | --- |
| Phenotype | Number | Genotype | Number |
| Long hypocotyl | 11 | *koy2/koy2* | 11 (100%) |
|  |  | WT/*koy2* | 0 |
|  |  | WT/WT | 0 |
| Short hypocotyl | 47 | *koy2/koy2* | 0 |
|  |  | WT/*koy2* | 35 (60.3%) |
|  |  | WT/WT | 12 (20.7%) |
